# Genome-wide mapping and prediction of plant architecture in a sorghum nested association mapping population

**DOI:** 10.1101/2020.01.28.923540

**Authors:** Marcus O. Olatoye, Zhenbin Hu, Geoffrey P. Morris

## Abstract

Modifying plant architecture is often necessary for yield improvement and climate adaptation, but we lack understanding of the genotype-phenotype map for plant morphology in sorghum. Here, we use a nested association mapping (NAM) population that captures global allelic diversity of sorghum to characterize the genetics of leaf erectness, leaf width (at two stages), and stem diameter. Recombinant inbred lines (n = 2200) were phenotyped in multiple environments (35,200 observations) and joint linkage mapping was performed with ∼93,000 markers. Fifty-four QTL of small to large effect were identified for trait BLUPs (9–16 per trait) each explaining 0.4–4% of variation across the NAM population. While some of these QTL colocalize with sorghum homologs of grass genes [e.g. involved in hormone synthesis (maize *spi1*), floral transition (*SbCN8*), and transcriptional regulation of development (rice *Ideal plant architecture1*)], most QTL did not colocalize with an *a priori* candidate gene (82%). Genomic prediction accuracy was generally high in five-fold cross-validation (0.65–0.83), and varied from low to high in leave-one-family-out cross-validation (0.04–0.61). The findings provide a foundation to identify the molecular basis of architecture variation in sorghum and establish genomic-enabled breeding for improved plant architecture.

**Core ideas:**

1. Understanding the genetics of plant architecture could facilitate the development of crop ideotypes for yield and adaptation
2. The genetics of plant architecture traits was characterized in sorghum using multi-environment phenotyping in a global nested association mapping population
3. Fifty-five quantitative trait loci were identified; some colocalize with homologs of known developmental regulators but most do not
4. Genomic prediction accuracy was consistently high in five-fold cross-validation, but accuracy varied considerably in leave-one-family-out predictions

## Introduction

Modification of plant architecture during crop improvement has greatly contributed to global agricultural productivity during the last century (Khush, 2001; Duvick, 2005). Ideotype breeding involves the creation of a model plant with characteristics that facilitate efficient photosynthesis, growth, and yield (Donald, 1968; Messina et al., 2009). The Green Revolution ideotypes targeted in maize, rice, and wheat includes reduced height, erect leaves, thick stalks, large and semi-compact inflorescence (ear, panicle, or head) (Khush, 2001). Several plant architecture traits have been under investigation for ideotype breeding. Leaf erectness (the angle between the culm and leaf midrib) is thought to have contributed to increased grain yield in maize through adaptation of hybrids to high planting densities (Duvick, 2005; Hammer et al., 2009). Wide leaves may facilitate efficient solar radiation capture for increased plant productivity (Sarlikioti et al., 2011). Thick stems may increase stalk strength and improve standability (Kashiwagi et al., 2008).

Genetic variation for these architectural traits exists in cereals that can be utilized in crop improvement (Tian et al., 2011; Zhao et al., 2011; Jia et al., 2013). Variation in above-ground plant structures may be due to genetic differences in regulators of maintenance, determinacy, and structure in apical meristem (Pautler et al., 2013). The leaves and shoots are generated from the shoot apical meristem at the vegetative phase, while the inflorescence meristems differentiate after floral transition (Wang and Li, 2008). Some key regulatory genes underlying plant architecture in grasses are *liguleless* genes (*lg1*–*lg4*) (Moreno et al., 1997; Johnston et al., 2014), *brachytic2* (*br2*) (Multani et al., 2003), *drooping leaf* (*drl1/drl2*) (Strable et al., 2017), and *Ideal Plant Architecture1 (IPA1)* (Miura et al., 2010; Liu et al., 2019). Growth hormones such as auxins, gibberellins, and brassinosteroids also regulate plant architecture (Wang and Li, 2008). Notably, major plant architecture regulators often have pleiotropic effects on other traits such as inflorescence shape, yield components, and disease resistance that can enhance or diminish their utility for crop improvement (Ishii et al., 2013; Kim et al., 2018; Liu et al., 2019).

Understanding trait genetic architecture is essential to design effective breeding strategies (Bernardo, 2008). Genetic architecture can characterized by various quantitative trait loci (QTL) mapping approaches, each with strengths and weaknesses (Morrell et al., 2012). While most crop trait dissection has used biparental populations, the effectiveness of these studies were generally limited by small population size and low genetic diversity. Association mapping exploits wide genetic diversity, but is limited by population structure that causes spurious and synthetic associations (Korte and Farlow, 2013). Nested association mapping (NAM) reduces the confounding effect of historical population structure and phenotypic variation through crosses between founder lines and common founder in NAM (Yu et al., 2008). Thus, NAM is an efficient approach to characterize the genetic basis of complex traits, especially adaptive traits that may be confounded with population structure (Bouchet et al., 2017).

Sorghum is an important staple food crop in semi-arid and arid regions globally due to its resilience to challenging environmental conditions (Monk et al., 2014). Sorghum cultivation in temperate climates during the last century was made possible through conversion of tall photoperiod-sensitive tropical germplasm to semi-dwarf photoperiod-insensitive lines (Klein et al., 2008). Genetics of flowering time and plant height in global sorghum diversity has been well characterized (Morris et al., 2013; Thurber et al., 2013; Bouchet et al., 2017), but less so for vegetative architecture traits such as leaf morphology and stem diameter (Zhao et al., 2016). Natural variation for plant architecture has been well characterized in maize, a relative of sorghum in tribe Andropogoneae, and found to be controlled by loci small effect (a predominantly polygenic architecture) (Tian et al., 2011; Li et al., 2015). However, evolutionary genetic theory suggests that sorghum, as a predominantly selfing species (Barnaud et al., 2008), would be expected to have a greater proportion of moderate to large effect variants than maize, an outcrossing species (Griswold, 2006).

In this study we characterized the genetic basis (location of QTL, number of QTL, effect sizes) of vegetative morphology traits in sorghum using a large NAM population and tested the accuracy of genomic prediction. Our results suggest that natural variation for vegetative traits in sorghum is under the control of a few loci of moderate effect and many loci of small effect (i.e. variation has oligogenic and polygenic components). While some QTL colocalized with homologs of maize and rice candidate genes, most did not, suggesting that natural variation for vegetative morphology in sorghum may be due predominantly to genes not previously described in other cereals.

## Materials and Methods

### Plant materials and phenotypic evaluation

The sorghum NAM population consists of 2200 recombinant inbred lines (RILs) derived from a cross between a common founder RTx430 (an important U.S. pollinator parent line) and 10 other diverse founders chosen to capture a wide genetic and morphological diversity (Table S1) (Bouchet et al., 2017). The genotyping of the population and the genome-wide SNP data set were previously described (Bouchet et al., 2017; Hu et al., 2019). The population was phenotyped in multiple environments in Kansas, US (Table 1) for leaf erectness, leaf width, stem diameter, flowering time and height. Leaf erectness (LFE) was defined as the angle between the pre-flag leaf midrib and a line perpendicular to culm at the point of pre-flag leaf attachment, and measured using a barcode reader and barcoded protractor (Figure S1). Leaf erectness was measured from three plants per plot. The leaf width was measured as the width of the leaf at the widest point on both pre-flag leaf and the fourth leaf from the flag leaf on three plants per plot (Figure S1). Measurements were taken using a barcode-scanning ruler. Stem diameter was measured as the diameter of the stem at the second internode above the ground, on three plants per plot, using a digital caliper. Flowering time was scored as the number of days from planting to the day in which 50% of the individuals in a plot are in anthesis. Inflorescence architecture phenotypes were previously described (Olatoye et al., 2019).

**Table 1:** Multi-environmental phenotyping of sorghum nested association mapping population.

| Location | Climate | Year | Annual<br>Precipitation<br>(mm)<sup>a</sup> | Environment Code | Traits<sup>b</sup> |
| --- | --- | --- | --- | --- | --- |
| Manhattan, KS | Humid<br>Continental | 2014 | 698 | MN14 | LET, PLW,<br>VLW, STM,<br>FLT |
| Hays, KS | Semi-Arid | 2014 | 639 | HA14 | STM, HGT |
| Hays, KS | Semi-Arid | 2015 | 513 | HI15 | LET, VLW,<br>PLW, HGT |

<sup>a</sup> Precipitation from October of the preceding year to October of the given year.
<sup>b</sup> Phenotypic traits evaluated are Leaf erectness (LET), pre-flag leaf width (PLW), vegetative leaf width (VLW), stem diameter (STM), height (HGT), and flowering time (FLT).

### Phenotypic data analysis

Analysis of phenotypic data collected was performed using R using LME4 package (Bates et al., 2015). The phenotypic mean of each RIL across environments was estimated. Genomic heritability for each trait was calculated by dividing genetic variance component estimate by the sum of genetic variance and residual variance component obtained using *mixed.solve* function in the rrBLUP R package (Endelman, 2011). Pairwise Pearson correlation among traits were estimated using the residuals derived from fitting a linear model for family and trait phenotypic means:

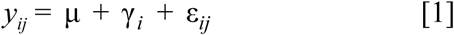

where *y*_*ij*_ is the phenotype, γ_*i*_ is the term for the NAM families and ε_*ij*_ is the residual term. The BLUPs (best linear unbiased predictors) for each phenotype was estimated by fitting the term for phenotyper (information on people that measured trait on the field) as fixed effect, while RIL, environment, RIL by environment interaction, and RIL by phenotype terms as random.

### Joint linkage mapping

Joint linkage mapping was performed using single environment phenotypes and trait BLUPs using the stepwise regression approach implemented in TASSEL 5.0 (Glaubitz et al., 2014). The approach is based on forward inclusion and backward elimination stepwise methods. The entry and exit limit of the forward and backward stepwise regressions were set at 0.001. The threshold cutoff was set at 1.8 E-6 based on 100 permutations. The genotypic data used for this analysis have been previously described (Hu et al., 2019). The JL model was specified as:

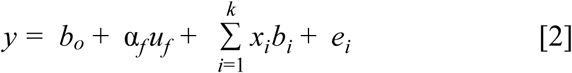

where *b*_0_ is the intercept, *u*_*f*_ is the effect of the family of the founder *f* obtained in the cross with the common parent (RTx430), α_*f*_ is the coefficient matrix relating *u*_*f*_ to **y**, *b*_*i*_ is the effect of the i^th^ identified locus in the model, *x*_*i*_ is the incidence vector that relates *b*_*i*_ to **y** and *k* is the number of significant QTL in the final model.

### Estimation of QTL effect size and allele frequency

The allele frequency of the QTL was estimated using code from http://evachan.org/calc_snp_stats.R. While the proportion of phenotypic variation explained (PVE) by each marker was estimated by fitting a regression model with terms for family and QTL nested within family as fixed effects (Würschum et al., 2012):

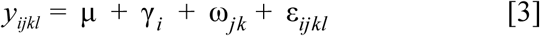

where *y*_*ijkl*_ is the phenotype, γ_i_ is the family term, ω_*jk*_ is the term for QTL nested within family, and ε_*ijk*_ is the residual. The sum of squares of marker nested within family divided by sum of squares total gave the proportion of variance explained by the detected QTL. The additive effect size of the QTL in the population was estimated as the average of the difference between the mean phenotypic values associated with the two-allele class of the QTL. The additive effect size of each QTL was estimated relative to the common parent RTx430.

### *Comparison of QTL regions with* a priori *candidate genes*

A list of *a priori* candidate genes was developed for leaf morphology, stem diameter, plant height, and flowering time based on homology with genes underlying these traits in other cereals (Supplementary File 1). The list included 14 sorghum genes and 107 sorghum homologs of maize and rice genes, which were identified based on “Protein Homologs” list for each maize or rice gene on Phytozome (Goodstein et al., 2012). The sorghum homolog with the highest similarity to the known gene was considered the putative ortholog and other homologs were considered putative paralogs. A custom R script was used to search for *a priori* genes that colocalized with the QTL within a window of 250 kb, the average distance until linkage disequilibrium decays to *r* < 0.1 in the NAM population (Hu et al., 2019).

### Genomic predictions

Genomic predictions were performed using the *mixed.solve* function in the ridge regression best linear unbiased prediction (rrBLUP) package in R (Endelman, 2011). The prediction models is as follows:

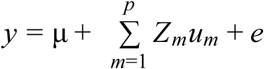

where **y** is the vector (*n* x 1) of observations, μ is the vector of the general mean, *p* is the marker number (*p* > *n*), Z_*m*_ is the *m*^th^ column of the design matrix **Z**, *u*_*m*_ is the genetic effect associated with *m*th marker, *p* is the number of markers, and *e* is the residual. Genomic prediction was cross-validated using two approaches. First, the “leave-one-family-out” prediction approach was performed by removing each family’s genotypic and phenotypic data and using remaining nine families to predict the given family. A five fold cross validation approach was used where the entire NAM population was divided into *n* = 5 equal groups, then *n* - 1 groups at a time were used to train the model to predict the remaining one. The process was then repeated for 100 cycles. For the five fold cross validation, two types of response variables were used in the genomic prediction model. First, the phenotypic mean across environments (no-family-effect), and second the residuals from the regression of trait values and family effect (family-effect).

## Results

### Phenotypic variation for plant architecture traits

Phenotypic observations were made on 2200 RILs in ten NAM families in three environments (year-by-location), resulting in 35,200 data points for vegetative traits. Phenotypic distribution of vegetative traits varied substantially among families (Figure 1; Table 2). Mean PLW ranged from 51 mm in the SC1103 family to 78 mm in the Macia family; mean VLW ranged from 62 mm in the SC1103 family to 97 mm in the Macia family; mean LFE ranged from 42 deg in the Ajabsido family to 66 deg in the SC1103 family; and mean STD ranged from 14 mm in the SC1103 family to 24 mm in the SC35 family. Phenotypic distributions were predominantly symmetrical, both with given families and with respect to the parent phenotype (Figure 1). In a few cases, RIL families had asymmetrical distributions (with respect to parent phenotypes) for some traits (e.g. LFE in the Ajabsido family, PLW in the SC1103 family).

**Table 2:** Summary of phenotypic and genotypic variation.

| **Trait<sup>a</sup>** | **Range** | **Mean** | **$h^2$<sup>b</sup>** | **Range of PVE (%)**<br>**for QTL<sup>c</sup>** | **Range of AES**<br>**for QTL<sup>d</sup>** |
| --- | --- | --- | --- | --- | --- |
| HGT (cm) | 53–270 | 109 | 0.53 | 0.2–14.7 | -12–7 |
| FLT (days) | 45–101 | 64 | 0.56 | 0.3–9.3 | -4–4 |
| STM (mm) | 10–34 | 19 | 0.43 | 0.4–1.6 | -2–3 |
| LFE (deg) | 13–85 | 56 | 0.36 | 0.6–4.1 | -3–7 |
| PLW (mm) | 33–112 | 68 | 0.31 | 0.4–2.6 | -6–6 |
| VLW (mm) | 45–131 | 86 | 0.25 | 0.5–1.2 | -5–8 |

<sup>a</sup> Preflag leaf width (PLW), vegetative leaf width VLW, leaf erectness (LFE), stem diameter (STD), plant height (HGT), and days to flowering time (FLT)
<sup>b</sup> Genomic heritability ( $h^2$ )
<sup>c</sup> Phenotypic variation explained
<sup>d</sup> Additive effect size

**Figure 1:**
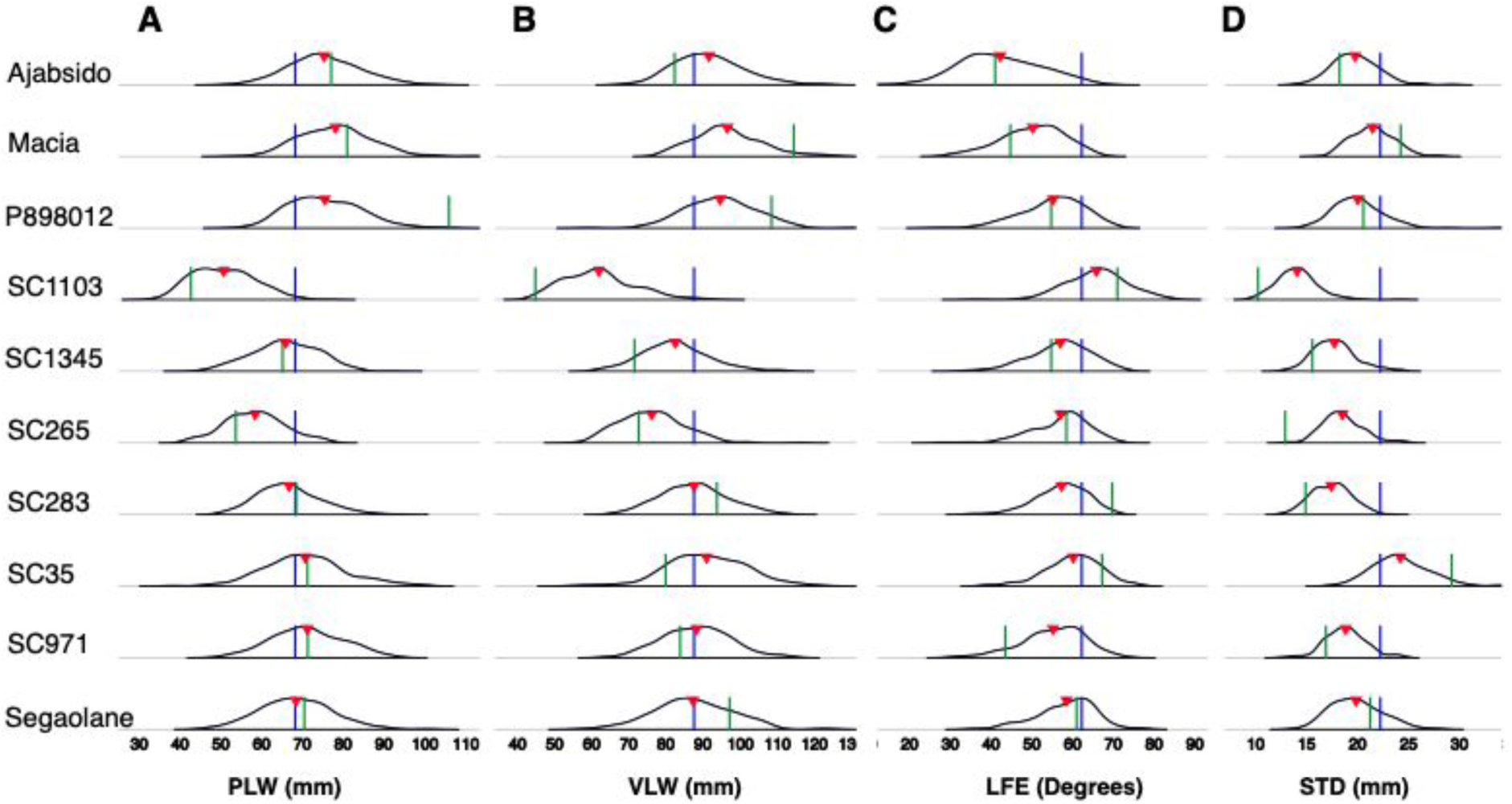
Phenotypic distribution of traits within sorghum nested association mapping (NAM) families. Density plots showing phenotypic distributions for (A) preflag leaf width (PLW), (B) vegetative leaf width (VLW; fourth leaf below the flag leaf,) (C) leaf erectness (LFE) of the preflag leaf, and (D) stem diameter (STD). Triangles indicate the mean trait value in each family, green lines indicates the trait value of the diverse founder for each family, and blue lines indicate the trait value of the common founder.

The relationship among pairs of traits, estimated as pairwise phenotypic correlations, differed substantially among trait pairs (Figure 2). LET had a weak negative correlation (*r* = -0.10; *P*-value < 0.01) with height and positive correlation with FLT (*r* = 0.19; *P*-value < 0.01) and STD (*r* = 0.16; *P*-value < 0.01), respectively. PLW was positively correlated with VLW (*r* = 0.46; *P*-value < 0.01), as expected, and had the same positive relationship with both STD and FLT (*r* = 0.20; *P*-value < 0.01). STD was positively correlated with FLT (*r* = 0.46; *P*-value < 0.01) and slightly negatively correlated to HGT (*r* = -0.09; *P*-value < 0.01).

**Figure 2:**
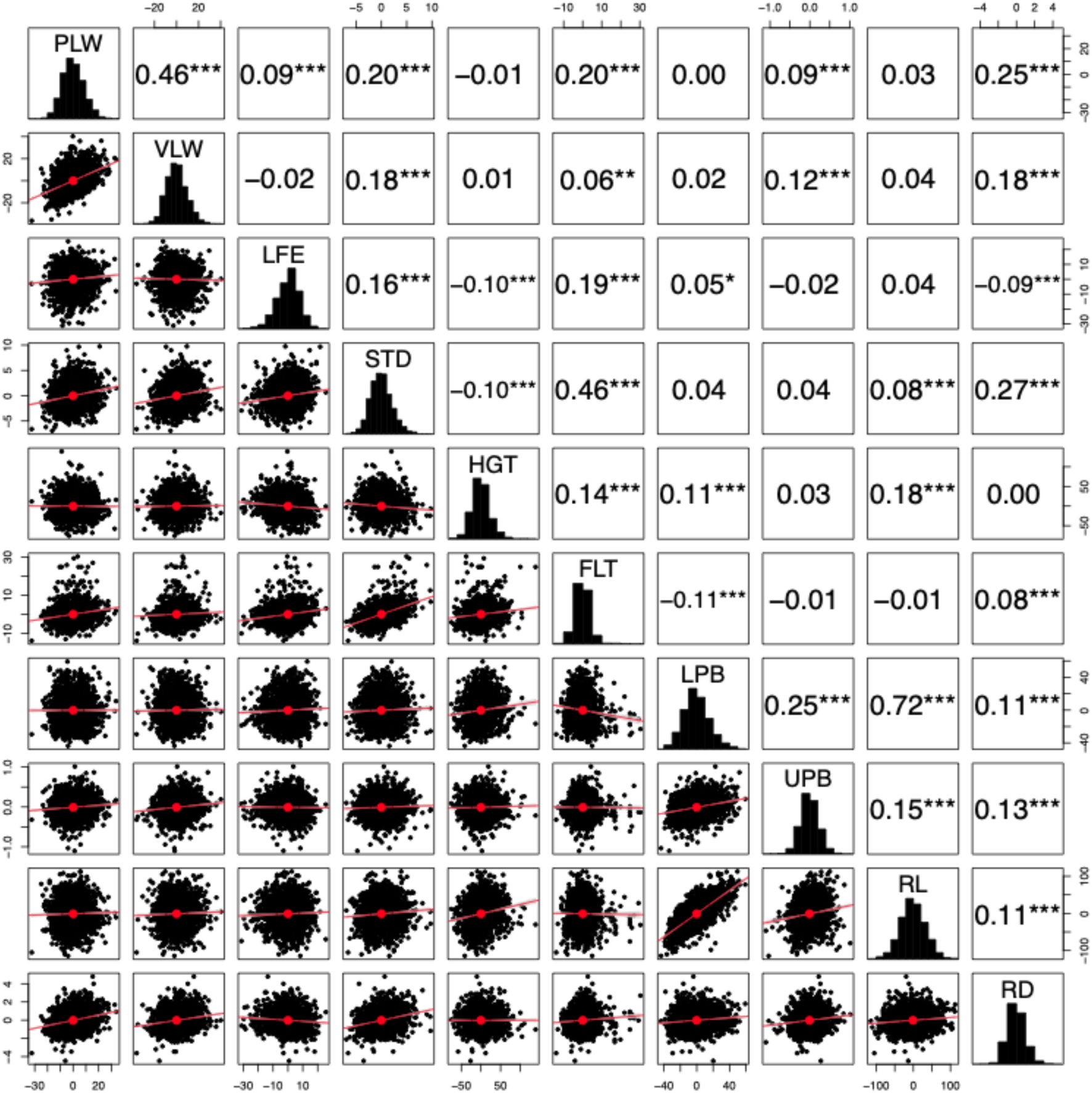
Relationships among plant architecture traits. Pairwise Pearson correlation among traits (preflag leaf width [PLW (mm)], vegetative leaf width [VLW (mm)], leaf erectness [LFE (degrees)], stem diameter [STD (mm)], plant height [HGT (cm)], flowering time [FLT (days)], lower primary branch [LPB (mm)], upper primary branch [UPB (mm)], rachis length [RL (mm)], and rachis diameter [RD (mm)]) after accounting for family effect (correlation of residuals of a linear model with a fixed family term). The correlation values shown are significant at *** *P*-value < 0.001, ** *P*-value < 0.01, and * *P*-value < 0.05.

### Identified QTL and their effect sizes

Joint linkage analysis was performed to characterize the genetic architecture of the four vegetative morphology traits (Table 2–3; Figure 3). Significant associations were observed for all traits, both in single environments and for trait BLUP across environments, with 60 QTL across all traits in single environment analysis (Supplementary File 2), and 54 QTL across all traits for BLUPs (Table 3). For trait BLUP associations, 16 SNPs explained 19% of LFE variation, 15 SNPs explained 15% of PLW variation, 9 SNPs explained 9% of VLW variation, and 14 SNPs explained 13% of STD variation. Allele frequencies of the QTL range from 0.05 to 0.45 for LFE, 0.05 to 0.39 for PLW, 0.10 to 0.33 for VLW, and 0.05 to 0.48 for STD. In general, the associated loci were of small to moderate effect when considering the proportion of variance explained across the entire NAM population. Among all traits, SNP S7_60258739, associated with LFE, explained the largest proportion of variation (4%). Among the 54 trait-associated SNPs, 50% of them (27 SNPs) explained less than 1% of phenotypic variation for the given trait.

**Table 3:**
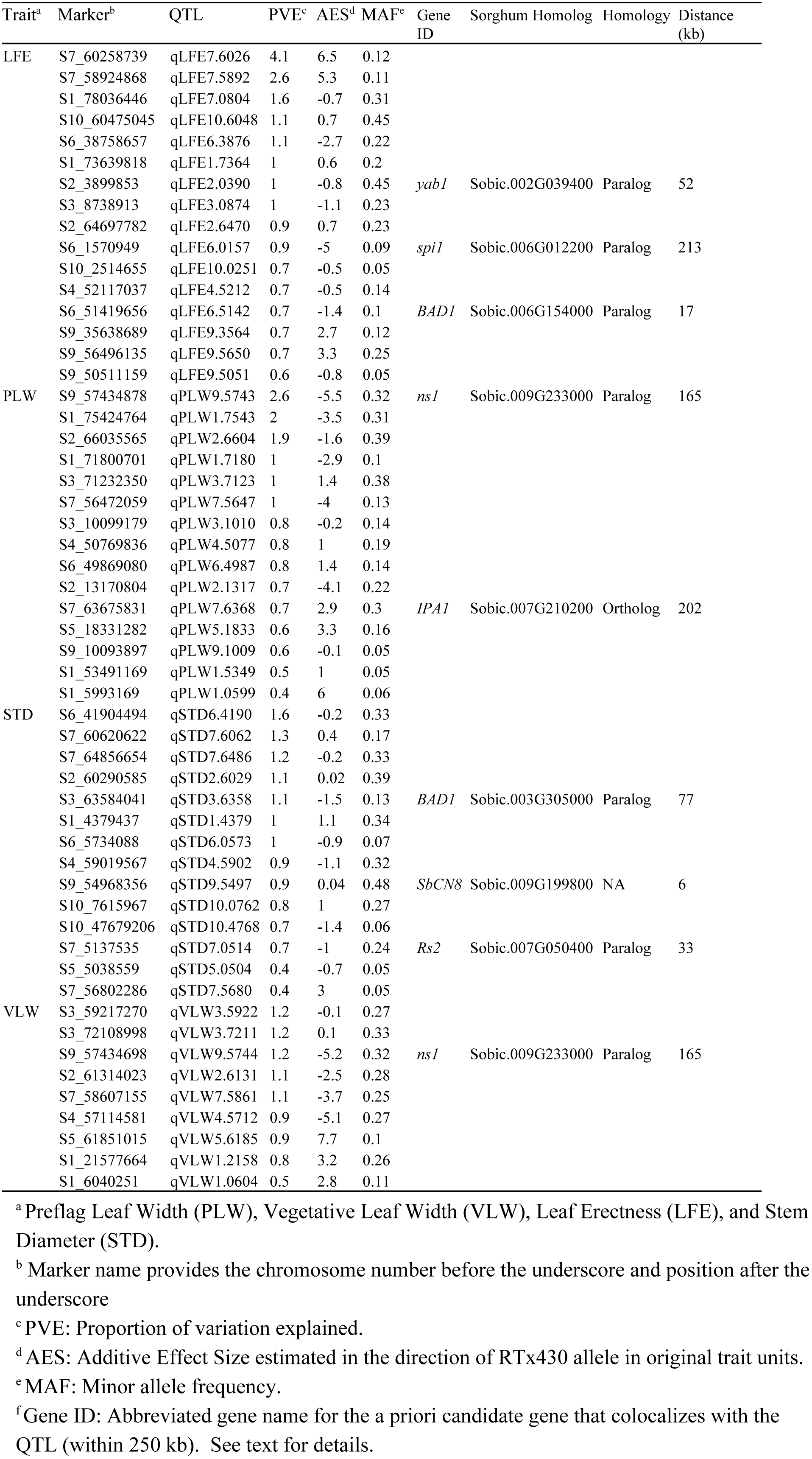
Markers associated with trait BLUPs in the NAM population.

**Figure 3:**
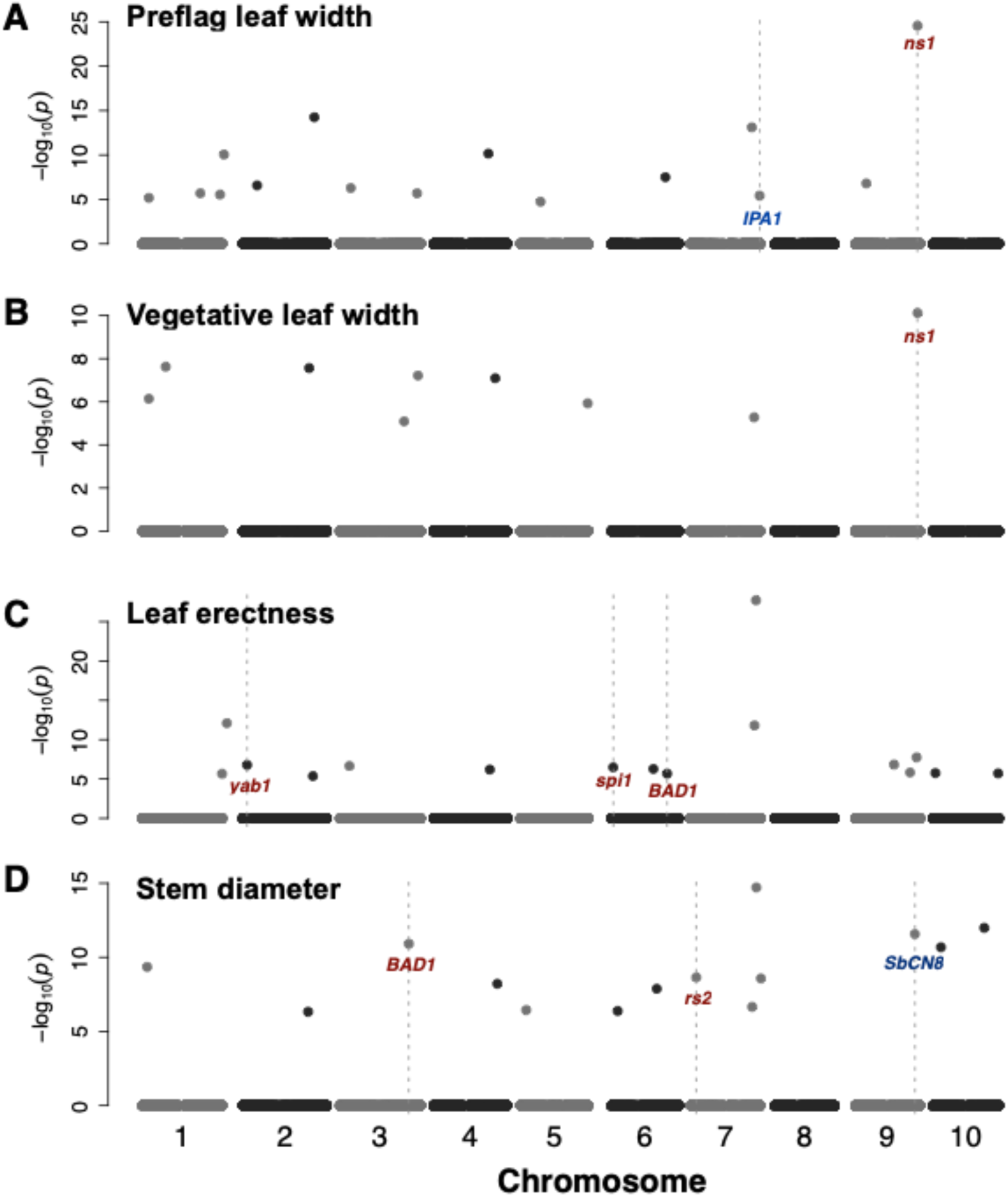
QTL mapping for vegetative morphology using joint linkage model. Genomic location of associations with best linear unbiased predictors of (A) preflag leaf width, (B) vegetative leaf width, (C) leaf erectness, and (D) stem diameter. *A priori* candidate genes that colocalize within 250 kb of QTL are noted, with putative orthologs of gene of interest in blue or putative paralogs in brown.

### QTL colocalized with plant and inflorescence morphology genes

Some of the QTL identified colocalized with *a priori* candidate genes (9 of 54; Figure 3; Table 3). For STD, a significant SNP (S7_59787744); PVE = 2%; MAF = 0.18) colocalized with two *a priori* candidate genes: the sorghum auxin efflux transporter gene (Sobic.007G163800; *Dwarf3*) at a distance of 34 kb and a sorghum paralog (Sobic.007G163200; *YUCCA5*) of the maize *sparse inflorescence1* (*spi1*) at a distance of 35 kb. A STD-associated SNP (S9_54968356; PVE = 1%; MAF = 0.48) colocalized 6 kb from *SbCN8* (Sobic.009G199800) a previously characterized sorghum flowering time regulator. For LET, a significant SNP (S7_63865966; PVE = 2%; MAF = 0.19) was 12 kb from the putative sorghum ortholog (Sobic.007G210200) of rice *Ideal Plant Architecture1* gene (*IPA1*). Across all vegetative trait QTL (for BLUPs and single environment mapping) (n = 114), 94 QTL (82%) did not colocalize with an *a priori* candidate gene, within the 250 kb LD window considered in this study.

### Genome-wide prediction accuracies

To assess the potential for genome-wide prediction of plant architecture, we cross-validated genomic BLUPs of eight traits using two approaches (Figure 4). We contrasted genomic prediction accuracy for vegetative architecture traits with accuracies for inflorescence architecture traits. With the five-fold cross-validation approach, moderate to high prediction accuracies were observed for all the traits when the residuals obtained from regression of phenotypes on family (family-effect model, FE; *r* = ∼0.2–0.6), When raw phenotypic data were used as the response variable in the genomic prediction model (no-family-effect model, NFE) accuracies were higher (*r* = ∼0.6–0.8). Mean prediction accuracy (across five folds and 100 cycles) for the FE model ranged from 0.25 in VLW to 0.41 in STD, and 0.42 in RD to 0.60 in RL. Mean prediction accuracy for the NFE model ranged from 0.69 in LFE to 0.81 in STD and 0.65 in UPB to 0.83 in RD (Figure 4A-B). For the leave-one-family-out approach, prediction accuracies were low to moderate, ranging from 0.10 to 0.6 across all traits (Figure 4C-D). For the leave-one-family-out approach, prediction accuracies were higher when certain families were used as validation sets (e.g. Ajabsido, P898012, SC35, and Segaolane). Inflorescence architecture traits were generally better predicted than vegetative architecture traits, other than RD, which was poorly predicted.

**Figure 4:**
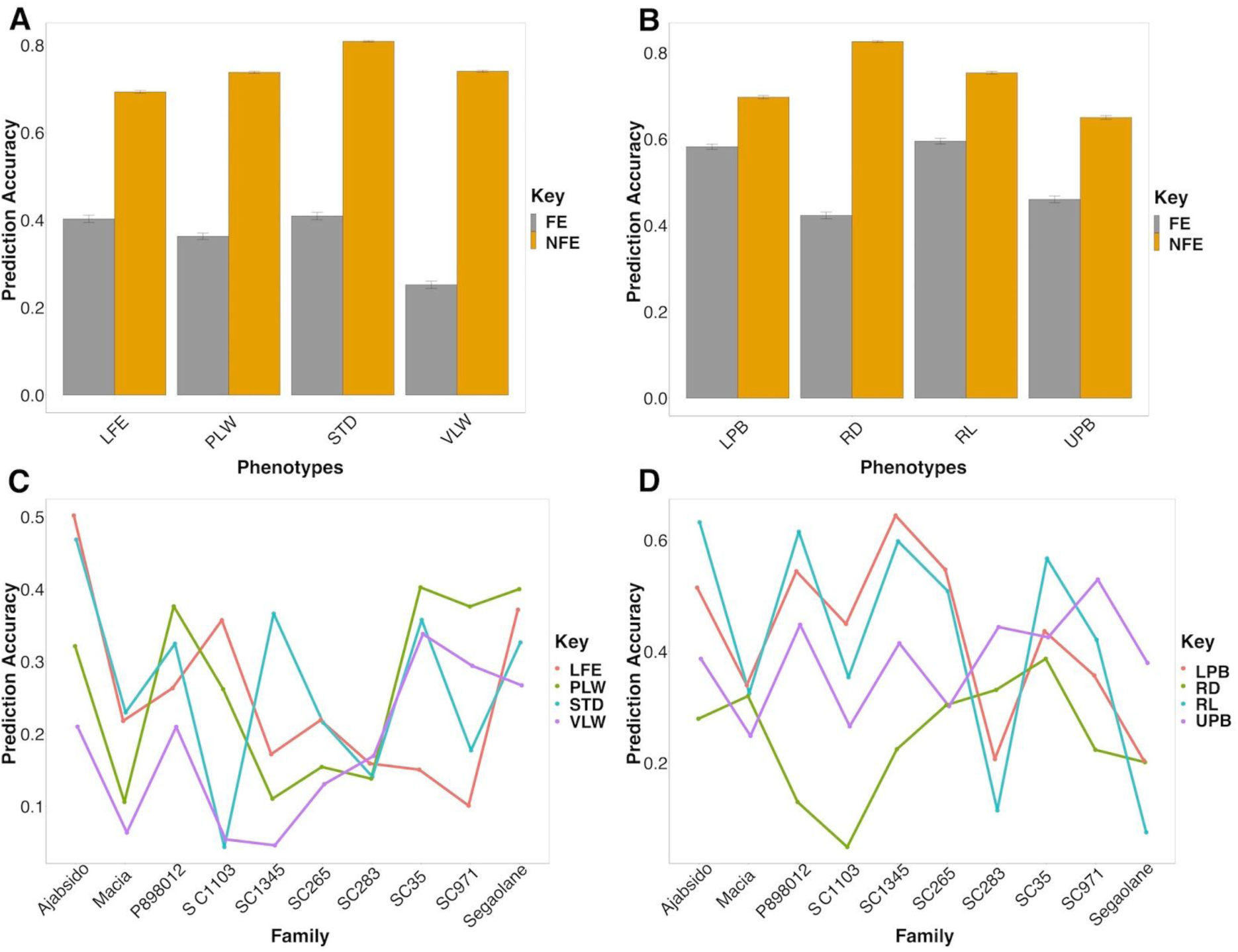
Genomic prediction accuracy for plant architecture traits. Upper panels are prediction accuracies from five-fold cross-validation for vegetative (A) and inflorescence (B) architecture traits. For “no family effect” (NFE) models, the response variables were the best linear unbiased predictors (BLUPs) of trait values. For the “family effect” (FE) models, the response variables were the residuals of trait value BLUPs, after a regression on family. Lower panels are prediction accuracies from leave-one-family-out approach, plotted by family for vegetative (C) and inflorescence (D) architecture traits. For all panels the prediction accuracies are Pearson correlations between observed trait value BLUPs and predicted trait values. Trait abbreviations: leaf erectness (LFE), preflag leaf width (PLW), stem diameter (STD), vegetative leaf width (VLW), lower primary branch (LPB), rachis diameter (RD), rachis length (RL), upper primary branch (UPB).

## Discussion

### The genetic architecture of plant architecture in sorghum

This study revealed that a few loci explaining moderate proportion of phenotypic variation vegetative trait are segregating in global sorghum diversity. All QTL associated with leaf traits were of small to moderate effect, explaining less than 4% of phenotypic variation. In maize, small effect loci (explaining < 4% of phenotypic variation) were found for both leaf erectness and leaf width (Tian et al., 2011). Mating system (outcrossing versus selfing) of species shapes the genetic architecture of complex traits, with selfing species expected to harbor a greater proportion of large effect QTL than outcrossers (Buckler et al., 2009; Tian et al., 2011). Our findings suggest that the relationship between mating system and QTL effect size may be trait specific, since leaf morphology traits QTL in both sorghum and maize NAM explain < 4% of phenotypic variation.

A previous study of a biparental mapping population identified a colocalized LFE and plant height QTL on chromosome 7 (Hart et al., 2001) which may correspond to the QTL we observe around 60 Mb on chromosome 7 (Table 3). Earlier QTL mapping of leaf erectness in sorghum using biparental populations identified an average of four QTL per population, with most QTL having an effect size of about 15% (Truong et al., 2015). However, the estimated effect size of these loci may have been inflated due to the Beavis effect in these small mapping populations (Xu, 2003). The allelic effect estimates in this study are expected to be more accurate due to the large number of RILs in the NAM population (i.e. 2200 RILs) (King and Long, 2017). A positive relationship was observed between heritability and the number of associated QTL since only the two low heritable leaf width traits were the ones with the least QTL number (compare Table 2 and Table 3).

Pleiotropy is particularly important for ideotype breeding, since selection on one trait may inadvertently change the phenotypic mean of other traits that are under shared genetic controls (Price and Langen, 1992). The modest negative correlation (*r* = -0.10) between HGT and LFE suggest that a short plants generally will have more erect leaves, which may facilitate adaptation for better light interception under high-density planting. The positive relationship between FLT and STM, is consistent with the expectation that late-flowering plants should have increased stem width due to the accumulation of biomass (Ashworth et al., 2016).

### Candidate genes for plant architecture variation

NAM provides an approach to investigate potential genes underlying natural variation. While fine mapping is needed to positively identify causal genes, a lack of colocalization of *a priori* candidate genes with NAM QTL with can provide evidence that the given genes do not underlie common variation. Given the evidence that *liguleless* genes condition natural variation for leaf angle in maize (Tian et al., 2011) we considered the hypothesis that *liguleless* homologs also underlie leaf angle variation in sorghum. However, no LFE QTL colocalized with homologs of *lg* genes (Table 3), suggesting that in contrast to maize *liguleless* genes do not underlie common natural variation for leaf angle in sorghum.

A few of the significant associations in this study were located near *a priori* candidate genes for both plant architecture, inflorescence architecture, and flowering time (Table 3). Some of these genes are known hormone transporters like the sorghum auxin transporter gene *Dw3* (Sobic.007G163800) which colocalized with both stem diameter and leaf erectness associated SNPs. Tall *Dw3* revertants have significantly more stem biomass than their short *dw3* counterparts (George-Jaeggli et al., 2013). Association studies in sorghum suggest a pleiotropic effect of *Dwarf3* on leaf erectness and inflorescence architecture (Brown et al., 2008). The colocalization of this gene alongside the negative correlation between stem diameter and plant height in this study suggests possible pleiotropic effect of the *Dw3* gene on stem diameter as previously observed for leaf erectness (Truong et al., 2015).

Plant development is dependent on transitions among several types of meristem and the network of genes that control them. A known regulator of vegetative-to-floral transition in sorghum, the florigen *SbCN8* (Yang et al., 2014), colocalized precisely (within 5 kb) with a STD-associated SNP on chromosome 9 (S9_54968356; qSTD9.549). The colocalization of a stem QTL with *SbCN8* would suggest that that *SbCN8* repression influences the control of vegetative growth, either via its effect on floral transition or by some other mechanism. However, its difficult to reconcile this hypothesis with the lack of a flowering time QTL at *SbCN8* in previous studies of the same NAM population (Bouchet et al., 2017; Hu et al., 2019).

The sorghum ortholog (Sobic.002G247800) of rice *Ideal Plant Architecture1* (*IPA1*, encoding an SPL transcription factor) colocalized with a common LFE-associated SNP (S7_63865966; PVE=1.5%, MAF=0.19). In rice, increase transcript accumulation of *IPA1* (also known as *OsSPL14* and *WFP)* resulted in reduced tiller number, stronger culms, and denser panicles (Jiao et al., 2010). Pleiotropy effects of the sorghum ortholog of *IPA1* with stem diameter in sorghum is also plausible since a significant STD-associated SNP colocalized with the sorghum ortholog of *IPA1* (200 kb away). Stem diameter is thought to be a component of lodging resistance and QTL for stem increased stem diameter have been used for the improvement of lodging resistance in rice (Kashiwagi et al., 2008). Another top QTL for LFE colocalized (60 kb) with a sorghum paralog (Sobic.002G039400) of maize *drooping leaf1* (*drl*) gene (Strable et al., 2017), with 35% similarity. The sorghum candidate gene is also ortholog of rice *YABBY1*, which creates a striking drooping leaf phenotype when cosuppressed by an overexpressed transgene (Dai et al., 2007).

The top QTL for both leaf width traits (VLW, PLW) colocalized with the WUSCHEL-related homeobox transcription factor Sobic.009G233000 (*SbWOX9*), an *a priori* candidate based on paralogy with maize *narrow sheath1* (*ns1*) gene (Nardmann et al., 2004). Note, *SbWOX9* is only a distant paralog of maize *ns1* (14% similar). However, *SbWOX9* is also the putative ortholog of rice *DWARF TILLER1* (Wang et al., 2014), which controls multiple aspects of plant architecture including internode width, suggesting this candidate may be worthy of further investigation.

Near isogenic line development, fine mapping, and/or molecular cloning of these candidate genes would be required to test the hypotheses that they contribute to variation in sorghum. Overall, however, our observation that many sorghum QTL do not colocalize with homologs of canonical maize and rice vegetative development regulators (e.g. most QTL in Fig. 3) suggests that much of the natural variation in vegetative morphology sorghum is due genes not previously described in cereals. This finding is consistent with recent molecular cloning studies, which have revealed that while some classical sorghum genes are orthologs of canonical genes known from model crops [e.g. *Tannin2* (Wu et al., 2019), *Maturity6* (Murphy et al., 2014)], many others are novel genes [e.g. *Dwarf1* (Yamaguchi et al., 2016), *Maturity2* (Casto et al., 2019), *Dry* (Zhang et al., 2018)].

### Prospects for genome-wide prediction of plant architecture

The high heritabilities and moderate prediction accuracies observed for each trait (Figure 4) suggest that genomic selection is possible for sorghum inflorescence morphology. This observation is despite the substantial oligogenic component which could make these traits less suited to genomic prediction models that assume polygenic variation (Bernardo, 2008). However, there is substantial variation in prediction accuracies obtained in the “leave-one-family-out” approach. This family-to-family variation may be a reflection of the differences in genetic architecture among NAM families of different botanical types. Prediction accuracies also varied among families in the maize NAM population (Peiffer et al., 2014), consistent with among-family differences in genetic architecture. The low prediction accuracies observed in some families (SC1103 for STD, RD, and VLW; SC1345 for VLW) suggests that these families may have variants or interactions which are not well-represented in the remaining NAM families. A better understanding of epistatic background effects may be needed to understand among-family variation in prediction accuracies (Blanc et al., 2006).

This genotype-phenotype map for plant architecture in sorghum has several potential uses in crop improvement. Trait-associated SNPs may be useful to develop molecular markers to facilitate simultaneous selection and improvement of multiple traits. However, attention has to be paid when selecting correlated traits that may have antagonistic effects on plant architecture or other aspects of agroclimatic adaptation (e.g. Figure 2). Finally, our study showed that the NAM population could be useful genomic selection training population for breeding population that share kinship with the NAM population founders.

## Abbreviations list

AES: Additive effect size
BLUP: Best linear unbiased predictor
FE: Family effect
FLT: Flowering time
HGT: Plant height
LFE: Leaf erectness
NAM: Nested association mapping
NFE: No family effect
PLW: Preflag leaf width
PVE: Proportion of variation explained
QTL: Quantitative trait loci
RIL: Recombinant inbred line
SNP: Single nucleotide polymorphism
STD: Stem diameter
VLW: Vegetative leaf width

## Supplemental Material statement

Supplementary File 1 contains the phenotypic data.

Supplementary File 2 contains (A) detailed descriptions of *a priori* genes (orthologs), (B) detailed descriptions of *a priori* genes (full homolog list), (C) QTL for all traits and environments, (D) QTL for vegetative traits BLUPs (E) QTL associated with vegetative traits’ in each environment, and (F) a table of *a priori* genes that colocalized with QTL.

## Data Availability statement

Genotype data are previously published and available at Dryad Digital Repository (doi:10.5061/dryad.gm073). Phenotype data are available in Supplementary File 1. Seed for the NAM population is available from the US National Plant Germplasm System.

## Author Contributions

MOO and GPM conceived and designed the study. MOO and ZH collected the data. MOO analyzed the data. MOO and GPM wrote the manuscript. All authors edited and approved the manuscript.

## Acknowledgements

This study is contribution 20-187-J from the Kansas Agricultural Experiment Station.

## Supplementary Figures and Tables

**Figure S1:**
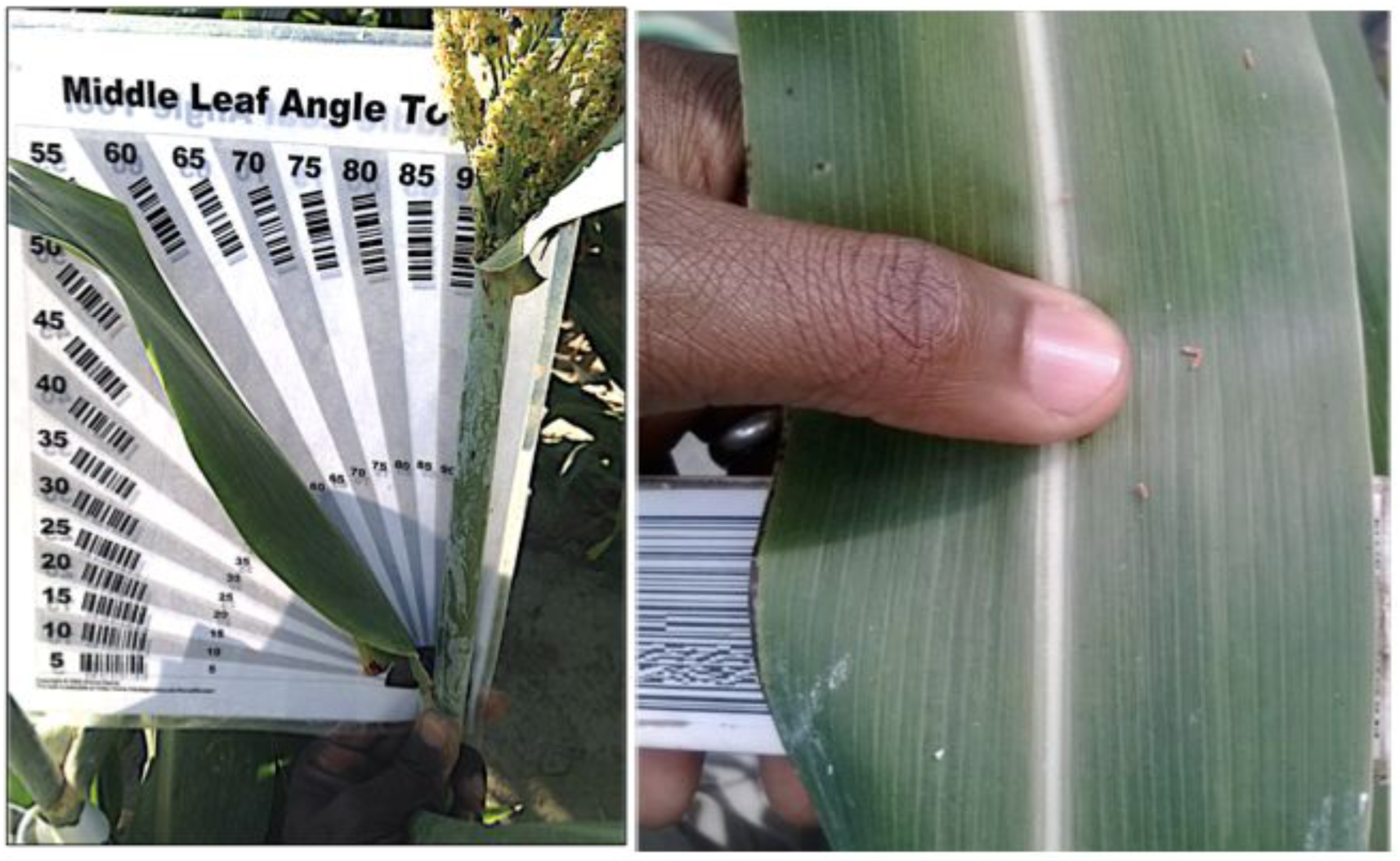
Phenotypic variation in leaf morphology. Leaf erectness and leaf width measurements. A barcode protractor and barcode ruler were both used for the measurements of these traits to ensure accurate and fast data acquisition.

**Table S1:**
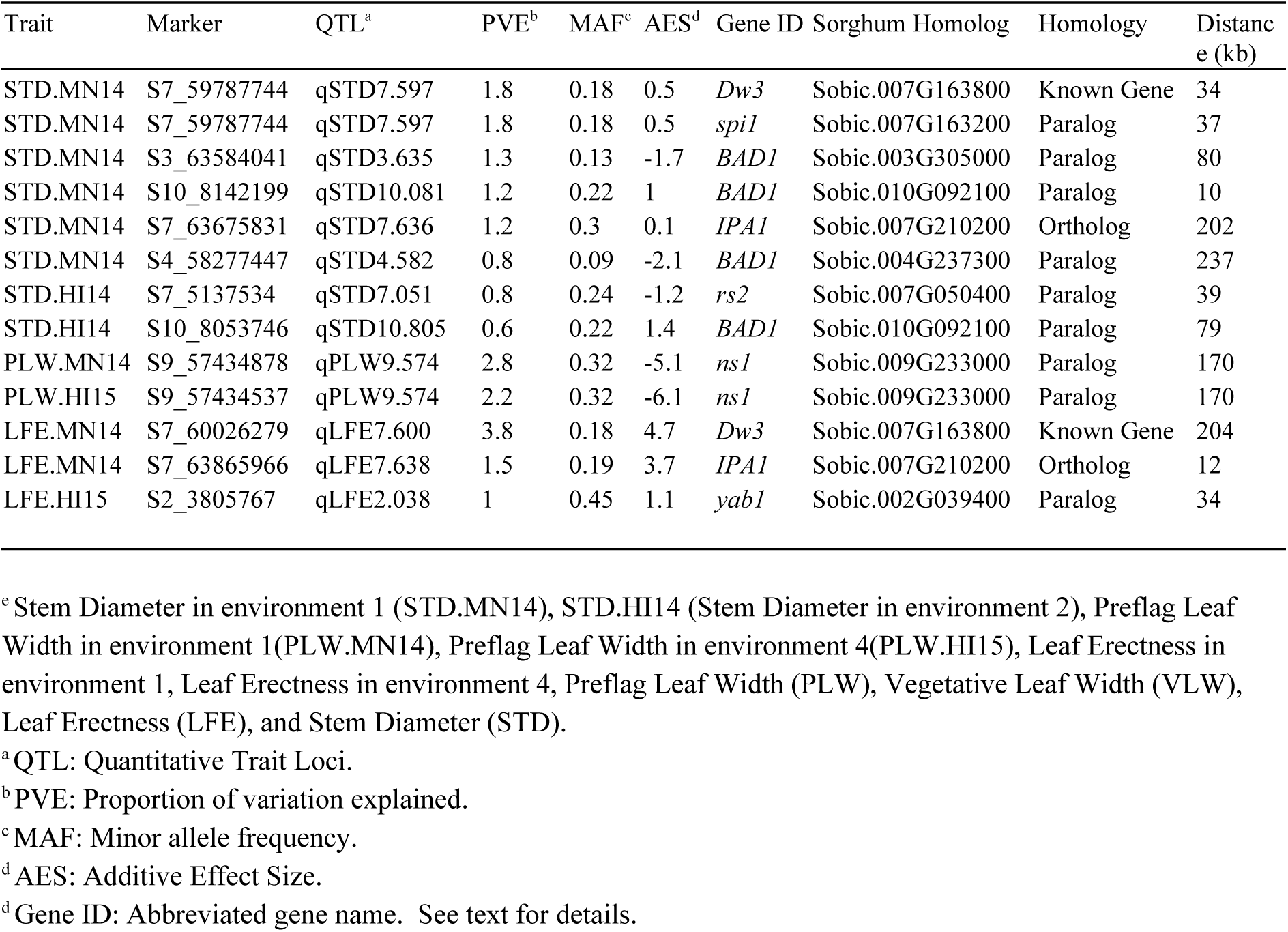
Joint linkage mapping significant markers that colocalize with *a priori* candidate genes within 250 kb.

## References

Ashworth, M.B., M.J. Walsh, K.C. Flower, M.M. Vila-Aiub, and S.B. Powles. 2016. Directional selection for flowering time leads to adaptive evolution in Raphanus raphanistrum (Wild radish). Evol. Appl. 9:619–629. doi: 10.1111/eva.12350

Barnaud, A., G. Trigueros, D. McKey, and H.I. Joly. 2008. High outcrossing rates in fields with mixed sorghum landraces: how are landraces maintained?. Heredity 101:445. doi: 10.1038/hdy.2008.77

Bernardo, R. 2008. Molecular markers and selection for complex traits in plants: Learning from the last 20 years. Crop Sci. 48:1649. doi: 10.2135/cropsci2008.03.0131

Blanc, G., A. Charcosset, B. Mangin, A. Gallais, and L. Moreau. 2006. Connected populations for detecting quantitative trait loci and testing for epistasis: an application in maize. Theor. Appl. Genet. 113:206–224. doi: 10.1007/s00122-006-0287-1

Bouchet, S., M.O. Olatoye, S.R. Marla, R. Perumal, T. Tesso, J. Yu, M. Tuinstra, and G.P. Morris. 2017. Increased power to dissect adaptive traits in global sorghum diversity using a nested association mapping population. Genetics 206:573–585. doi: 10.1534/genetics.116.198499

Buckler, E.S., J.B. Holland, P.J. Bradbury, C.B. Acharya, P.J. Brown, C. Browne, E. Ersoz, S. Flint-Garcia, A. Garcia, J.C. Glaubitz, M.M. Goodman, C. Harjes, K. Guill, D.E. Kroon, S. Larsson, N.K. Lepak, H. Li, S.E. Mitchell, G. Pressoir, J.A. Peiffer, M.O. Rosas, T.R. Rocheford, M.C. Romay, S. Romero, S. Salvo, H.S. Villeda, H.S.D. Silva, Q. Sun, F. Tian, N. Upadyayula, D. Ware, H. Yates, J. Yu, Z. Zhang, S. Kresovich, and M.D. McMullen. 2009. The genetic architecture of maize flowering time. Science 325:714–718. doi: 10.1126/science.1174276

Casto, A.L., A.J. Mattison, S.N. Olson, M. Thakran, W.L. Rooney, and J.E. Mullet. 2019. Maturity2, a novel regulator of flowering time in Sorghum bicolor, increases expression of SbPRR37 and SbCO in long days delaying flowering. PLOS ONE 14:e0212154. doi: 10.1371/journal.pone.0212154

Dai, M., Y. Zhao, Q. Ma, Y. Hu, P. Hedden, Q. Zhang, and D.-X. Zhou. 2007. The rice YABBY1 gene is involved in the feedback regulation of gibberellin metabolism. Plant Physiol. 144:121–133. doi: 10.1104/pp.107.096586

Donald, C.M. 1968. The breeding of crop ideotypes. Euphytica 17:385–403

Duvick, D.N. 2005. Genetic progress in yield of United States maize (Zea mays L.). Maydica 50:193

Endelman, J.B. 2011. Ridge regression and other kernels for genomic selection with R package rrBLUP. Plant Genome 4:250. doi: 10.3835/plantgenome2011.08.0024

George-Jaeggli, B., D.R. Jordan, E.J. van Oosterom, I.J. Broad, and G.L. Hammer. 2013. Sorghum dwarfing genes can affect radiation capture and radiation use efficiency. Field Crops Res. 149:283–290. doi: 10.1016/j.fcr.2013.05.005

Glaubitz, J.C., T.M. Casstevens, F. Lu, J. Harriman, R.J. Elshire, Q. Sun, and E.S. Buckler. 2014. TASSEL-GBS: A high capacity genotyping by sequencing analysis pipeline. PLoS ONE 9:e90346. doi: 10.1371/journal.pone.0090346

Goodstein, D.M., S. Shu, R. Howson, R. Neupane, R.D. Hayes, J. Fazo, T. Mitros, W. Dirks, U. Hellsten, N. Putnam, and D.S. Rokhsar. 2012. Phytozome: a comparative platform for green plant genomics. Nucleic Acids Res. 40:D1178–D1186. doi: 10.1093/nar/gkr944

Griswold, C.K. 2006. Gene flow’s effect on the genetic architecture of a local adaptation and its consequences for QTL analyses. Heredity 96:6800822. doi: 10.1038/sj.hdy.6800822

Hammer, G.L., Z. Dong, G. McLean, A. Doherty, C. Messina, J. Schussler, C. Zinselmeier, S. Paszkiewicz, and M. Cooper. 2009. Can changes in canopy and/or root system architecture explain historical maize yield trends in the US corn belt?. Crop Sci. 49:299–312

Hart, G.E., K.F. Schertz, Y. Peng, and N.H. Syed. 2001. Genetic mapping of Sorghum bicolor (L.) Moench QTLs that control variation in tillering and other morphological characters. Theor. Appl. Genet. 103:1232–1242. doi: 10.1007/s001220100582

Hu, Z., M.O. Olatoye, S. Marla, and G.P. Morris. 2019. An integrated genotyping-by-sequencing polymorphism map for over 10,000 sorghum genotypes. Plant Genome 12:1–15. doi: 10.3835/plantgenome2018.06.0044

Jia, G., X. Huang, H. Zhi, Y. Zhao, Q. Zhao, W. Li, Y. Chai, L. Yang, K. Liu, H. Lu, C. Zhu, Y. Lu, C. Zhou, D. Fan, Q. Weng, Y. Guo, T. Huang, L. Zhang, T. Lu, Q. Feng, H. Hao, H. Liu, P. Lu, N. Zhang, Y. Li, E. Guo, S. Wang, S. Wang, J. Liu, W. Zhang, G. Chen, B. Zhang, W. Li, Y. Wang, H. Li, B. Zhao, J. Li, X. Diao, and B. Han. 2013. A haplotype map of genomic variations and genome-wide association studies of agronomic traits in foxtail millet (Setaria italica). Nat. Genet. 45:957–961. doi: 10.1038/ng.2673

Jiao, Y., Y. Wang, D. Xue, J. Wang, M. Yan, G. Liu, G. Dong, D. Zeng, Z. Lu, X. Zhu, Q. Qian, and J. Li. 2010. Regulation of OsSPL14 by OsmiR156 defines ideal plant architecture in rice. Nat. Genet. 42:541–544. doi: 10.1038/ng.591

Kashiwagi, T., E. Togawa, N. Hirotsu, and K. Ishimaru. 2008. Improvement of lodging resistance with QTLs for stem diameter in rice (Oryza sativa L.). Theor. Appl. Genet. 117:749–757. doi: 10.1007/s00122-008-0816-1

Khush, G.S. 2001. Green revolution: the way forward. Nat. Rev. Genet. 2:815. doi: 10.1038/35093585

King, E.G., and A.D. Long. 2017. The Beavis effect in next-generation mapping panels in Drosophila melanogaster. G3 Genes Genomes Genet. 7:1643–1652. doi: 10.1534/g3.117.041426

Klein, R.R., J.E. Mullet, D.R. Jordan, F.R. Miller, W.L. Rooney, M.A. Menz, C.D. Franks, and P.E. Klein. 2008. The effect of tropical sorghum conversion and inbred development on genome diversity as revealed by high-resolution genotyping. Crop Sci. 48:S–12. doi: 10.2135/cropsci2007.06.0319tpg

Korte, A., and A. Farlow. 2013. The advantages and limitations of trait analysis with GWAS: a review. Plant Methods 9:29. doi: 10.1186/1746-4811-9-29

Li, C., Y. Li, Y. Shi, Y. Song, D. Zhang, E.S. Buckler, Z. Zhang, T. Wang, and Y. Li. 2015. Genetic control of the leaf angle and leaf orientation value as revealed by ultra-high density maps in three connected maize populations. PLoS ONE 10. doi: 10.1371/journal.pone.0121624

Messina, C., G. Hammer, Z. Dong, D. Podlich, and M. Cooper. 2009. Modelling crop improvement in a G X E X M framework via gene-trait-phenotype relationships. Crop Physiol. Interfacing Genet. Improv. Agron. Neth. Elsevier 235–265

Monk, R., C. Franks, and J. Dahlberg. 2014. Sorghum. Crop Science Society of America.

Morrell, P.L., E.S. Buckler, and J. Ross-Ibarra. 2012. Crop genomics: advances and applications. Nat. Rev. Genet. 13:85–96. doi: 10.1038/nrg3097

Morris, G.P., P. Ramu, S.P. Deshpande, C.T. Hash, T. Shah, H.D. Upadhyaya, O. Riera-Lizarazu, P.J. Brown, C.B. Acharya, S.E. Mitchell, J. Harriman, J.C. Glaubitz, E.S. Buckler, and S. Kresovich. 2013. Population genomic and genome-wide association studies of agroclimatic traits in sorghum. Proc. Natl. Acad. Sci. 110:453–458. doi: 10.1073/pnas.1215985110

Murphy, R.L., D.T. Morishige, J.A. Brady, W.L. Rooney, S. Yang, P.E. Klein, and J.E. Mullet. 2014. Ghd7 (Ma6) represses sorghum flowering in long days: Ghd7 alleles enhance biomass accumulation and grain production. Plant Genome 7:1–10. doi: 10.3835/plantgenome2013.11.0040

Nardmann, J., J. Ji, W. Werr, and M.J. Scanlon. 2004. The maize duplicate genes narrow sheath1 and narrow sheath2 encode a conserved homeobox gene function in a lateral domain of shoot apical meristems. Development 131:2827–2839. doi: 10.1242/dev.01164

Olatoye, M.O., S.R. Marla, Z. Hu, S. Bouchet, R. Perumal, and G.P. Morris. 2019. Dissecting adaptive traits with nested association mapping: Genetic architecture of inflorescence morphology in sorghum. bioRxiv 748681. doi: 10.1101/748681

Pautler, M., W. Tanaka, H.-Y. Hirano, and D. Jackson. 2013. Grass meristems I: shoot apical meristem maintenance, axillary meristem determinacy and the floral transition. Plant Cell Physiol. 54:302–312. doi: 10.1093/pcp/pct025

Peiffer, J.A., M.C. Romay, M.A. Gore, S.A. Flint-Garcia, Z. Zhang, M.J. Millard, C.A.C. Gardner, M.D. McMullen, J.B. Holland, P.J. Bradbury, and E.S. Buckler. 2014. The genetic architecture of maize height. Genetics 196:1337–1356. doi: 10.1534/genetics.113.159152

Price, T., and T. Langen. 1992. Evolution of correlated characters. Trends Ecol. Evol. 7:307–310. doi: 10.1016/0169-5347(92)90229-5

Sarlikioti, V., P.H.B. de Visser, G.H. Buck-Sorlin, and L.F.M. Marcelis. 2011. How plant architecture affects light absorption and photosynthesis in tomato: towards an ideotype for plant architecture using a functional–structural plant model. Ann. Bot. 108:1065–1073. doi: 10.1093/aob/mcr221

Thurber, C.S., J.M. Ma, R.H. Higgins, and P.J. Brown. 2013. Retrospective genomic analysis of sorghum adaptation to temperate-zone grain production. Genome Biol. 14:R68. doi: 10.1186/gb-2013-14-6-r68

Tian, F., P.J. Bradbury, P.J. Brown, H. Hung, Q. Sun, S. Flint-Garcia, T.R. Rocheford, M.D. McMullen, J.B. Holland, and E.S. Buckler. 2011. Genome-wide association study of leaf architecture in the maize nested association mapping population. Nat. Genet. 43:159–162. doi: 10.1038/ng.746

Truong, S.K., R.F. McCormick, W.L. Rooney, and J.E. Mullet. 2015. Harnessing genetic variation in leaf angle to increase productivity of Sorghum bicolor. Genetics 201:1229–1238. doi: 10.1534/genetics.115.178608

Wang, W., G. Li, J. Zhao, H. Chu, W. Lin, D. Zhang, Z. Wang, and W. Liang. 2014. DWARF TILLER1, a WUSCHEL-related homeobox transcription factor, is required for tiller growth in rice. PLOS Genet. 10:e1004154. doi: 10.1371/journal.pgen.1004154

Wu, Y., T. Guo, Q. Mu, J. Wang, X. Li, Y. Wu, B. Tian, M.L. Wang, G. Bai, R. Perumal, H.N. Trick, S.R. Bean, I.M. Dweikat, M.R. Tuinstra, G. Morris, T.T. Tesso, J. Yu, and X. Li. 2019. Allelochemicals targeted to balance competing selections in African agroecosystems. Nat. Plants 1–8. doi: 10.1038/s41477-019-0563-0

Würschum, T., W. Liu, M. Gowda, H.P. Maurer, S. Fischer, A. Schechert, and J.C. Reif. 2012. Comparison of biometrical models for joint linkage association mapping. Heredity 108:332–340. doi: 10.1038/hdy.2011.78

Xu, S. 2003. Theoretical basis of the Beavis Effect. Genetics 165:2259–2268

Yamaguchi, M., H. Fujimoto, K. Hirano, S. Araki-Nakamura, K. Ohmae-Shinohara, A. Fujii, M. Tsunashima, X.J. Song, Y. Ito, R. Nagae, J. Wu, H. Mizuno, J. Yonemaru, T. Matsumoto, H. Kitano, M. Matsuoka, S. Kasuga, and T. Sazuka. 2016. Sorghum Dw1, an agronomically important gene for lodging resistance, encodes a novel protein involved in cell proliferation. Sci. Rep. 6:28366. doi: 10.1038/srep28366

Yang, S., R.L. Murphy, D.T. Morishige, P.E. Klein, W.L. Rooney, and J.E. Mullet. 2014. Sorghum phytochrome B inhibits flowering in long days by activating expression of SbPRR37 and SbGHD7, repressors of SbEHD1, SbCN8 and SbCN12. PLoS ONE 9:e105352. doi: 10.1371/journal.pone.0105352

Yu, J., J.B. Holland, M.D. McMullen, and E.S. Buckler. 2008. Genetic design and statistical power of nested association mapping in maize. Genetics 178:539–551. doi: 10.1534/genetics.107.074245

Zhang, L.-M., C.-Y. Leng, H. Luo, X.-Y. Wu, Z.-Q. Liu, Y.-M. Zhang, H. Zhang, Y. Xia, L. Shang, C.-M. Liu, D.-Y. Hao, Y.-H. Zhou, C.-C. Chu, H.-W. Cai, and H.-C. Jing. 2018. Sweet sorghum originated through selection of Dry, a plant-specific NAC transcription factor gene. Plant Cell 30:2286–2307. doi: 10.1105/tpc.18.00313

Zhao, J., M.B. Mantilla Perez, J. Hu, and M.G. Salas Fernandez. 2016. Genome-wide association study for nine plant architecture traits in sorghum. Plant Genome 9. doi: 10.3835/plantgenome2015.06.0044

Zhao, K., C.-W. Tung, G.C. Eizenga, M.H. Wright, M.L. Ali, A.H. Price, G.J. Norton, M.R. Islam, A. Reynolds, J. Mezey, A.M. McClung, C.D. Bustamante, and S.R. McCouch. 2011. Genome-wide association mapping reveals a rich genetic architecture of complex traits in Oryza sativa. Nat Commun 2:467. doi: 10.1038/ncomms1467

